# Gaze Behaviour Reveals Flexible Encoding of Competing Reach Goals Under Conditions of Target Uncertainty

**DOI:** 10.1101/2020.09.02.279414

**Authors:** Anouk J. de Brouwer, Michael J. Carter, Lauren C. Smail, Daniel M. Wolpert, Jason P. Gallivan, J. Randall Flanagan

## Abstract

In daily tasks, we are often confronted with competing potential targets and must select one to act on. It has been suggested that, prior to target selection, the human brain encodes the motor goals of multiple, potential targets. However, this view remains controversial and it has been argued that only a single motor goal is encoded, or that motor goals are only specified after target selection. To investigate this issue, we measured participants’ gaze behaviour while viewing two potential reach targets, one of which was cued after a preview period. We applied visuomotor rotations to dissociate each visual target location from its corresponding motor goal location; i.e., the location participants needed to aim their hand toward to bring the rotated cursor to the target. During the preview period, participants most often fixated both motor goals but also frequently fixated one, or neither, motor goal location. Further gaze analysis revealed that on trials in which both motor goals were fixated, both locations were held in memory simultaneously. These findings show that, at the level of single trials, the brain most often encodes multiple motor goals prior to target selection, but may also encode either one or no motor goals. This result may help reconcile a key debate concerning the specification of motor goals in cases of target uncertainty.

## Introduction

Preparing a reaching movement towards a visual target is thought to involve transforming the visual representation of the target into a motor representation, which constitutes the motor goal of the action (Crawford et al., 2004). In our everyday lives, we frequently encounter situations in which we must select between competing potential targets of action, as when choosing a particular coffee mug to reach for from a cupboard. A fundamental, and as yet unresolved, question is whether, prior to target selection, the brain specifies and maintains, in parallel, competing motor goals for different potential targets (Gallivan et al., 2018). According to the influential affordance competition hypothesis (Cisek, 2012, 2007; Cisek and Kalaska, 2010; Thura and Cisek, 2014), the brain specifies motor goals for competing options, in parallel, before deciding which one to execute (see also Klaes et al., 2011; Suriya-Arunroj and Gail, 2019). Although a number of behavioural and neurophysiological studies have expressed support for this hypothesis, alternative interpretations of the results of this work have been put forward; arguing that only a single motor goal is specified prior to target selection (Dekleva et al., 2018) or that the motor goal is specified only after target selection (Gallivan et al., 2018).

Support for the affordance competition hypothesis comes from single cell recording studies that have employed delayed reach tasks in which one of two potential targets is cued after a preview period. These studies have found that, prior to target selection, competing potential reach targets appear to be represented in parallel in brain areas thought to be directly involved in movement execution, including dorsal premotor cortex (Cisek and Kalaska, 2005; Coallier et al., 2015; Pastor-Bernier and Cisek, 2011) and the parietal reach region (Klaes et al., 2011). However, this parallel specification interpretation has recently been challenged by a study that simultaneously recorded activity from populations of neurons in dorsal premotor cortex during a delayed reach task with two potential targets (Dekleva et al., 2018). The authors of this study argued that the apparent parallel representation of competing targets is an artifact of averaging across trials, and that, at the level of single trials, neural population activity is more consistent with only a single potential target being represented. That is, they argued in favour of the hypothesis that the brain only encodes a single motor goal, and then revises this motor goal in favour of the other target if necessary; what they referred to as a stay-or-switch model.

Behavioural studies have sought to test the parallel specification hypothesis using ‘go-before-you-know’ tasks. In such tasks, participants are simultaneously presented with two or more potential reach targets and are required to immediately launch a reach movement towards these competing targets *before* knowing the final target location, which is cued after movement onset (Chapman et al., 2010; Gallivan et al., 2017, 2016b, 2011; Haith et al., 2016; Stewart et al., 2014, 2013; Wong and Haith, 2017). In these tasks, the initial reach is typically directed towards the midpoint of the potential targets, leading to the initial suggestion that the motor system rapidly forms a motor plan for each potential target and then executes an average of these plans (Chapman et al., 2010). However, it has been shown that launching movements in an intermediate spatial direction minimizes motor costs associated with corrective movements (Christopoulos et al., 2015; Christopoulos and Schrater, 2015; Hudson et al., 2007), and it has been argued that ‘averaging’ behaviour arises from executing a single movement that is optimized based on task constraints (Gallivan et al., 2018, 2017; Haith et al., 2015; Nashed et al., 2017; Wong and Haith, 2017). Using a delayed reach task, we provided behavioural evidence that motor goals of two potential targets are encoded during the preview period (Gallivan et al., 2015). However, it is possible that a stay-or-switch model could also account for the results of that study.

We recently examined gaze behaviour in a delayed reaching task with a single target under a visuomotor rotation (de Brouwer et al., 2018). We found that during the preview period, participants—in addition to fixating the visual target—reliably fixated the motor goal; i.e., an ‘aimpoint’, rotated away from the target, to which they subsequently directed their reaching movement (Rand and Rentsch, 2015; see also Rentsch and Rand, 2014). Here we employed a variant of this task with two potential targets to assess, using gaze behaviour, whether people specify a single motor goal or multiple motor goals in single reach trials.

We show that participants frequently fixated, and retained in memory, the motor goal locations of both potential targets, providing support for the parallel specification hypothesis. However, we also find that participants often fixated only one of the two motor goals, as predicted by the stay-or-switch model, or neither motor goal, suggesting that motor goal specification occurred after target selection. Individual participants exhibited multiple fixation strategies suggesting that individuals can flexibly alternate between different modes of motor goal encoding. These results may serve to reconcile seemingly disparate findings from previous studies that have assumed that only one encoding strategy is in operation, rather than a mixture of strategies.

## Results

To assess the encoding of motor goals prior to target selection, we had participants perform a center-out reaching task in which they moved a cursor to visual targets presented on a vertical monitor. This was accomplished by sliding a hand-held stylus across a drawing tablet without vision of the hand, with forward and rightward stylus motion corresponding to upward and rightward cursor motion under baseline conditions. In each trial, one or two targets were presented on a visible ring composed of 60 small circles, with blue targets always appearing at one of four locations on the left half of the ring, and red targets always appearing at one of four locations on the right side of the ring (Fig. 1a). In 1-target reach trials, either a blue or a red target was displayed for a 2-s preview period, and in 2-target reach trials, one blue target and one red target were displayed for a 4-s preview phase. During the preview phase, all targets were presented as open circles. At the end of the preview phase, either the single target or one or the two targets was ‘filled in’, cueing the participant to reach to that target. Participants were instructed to make rapid movements such that the cursor ‘sliced’ through the target, thereby minimizing corrective hand actions.

**Figure 1.**
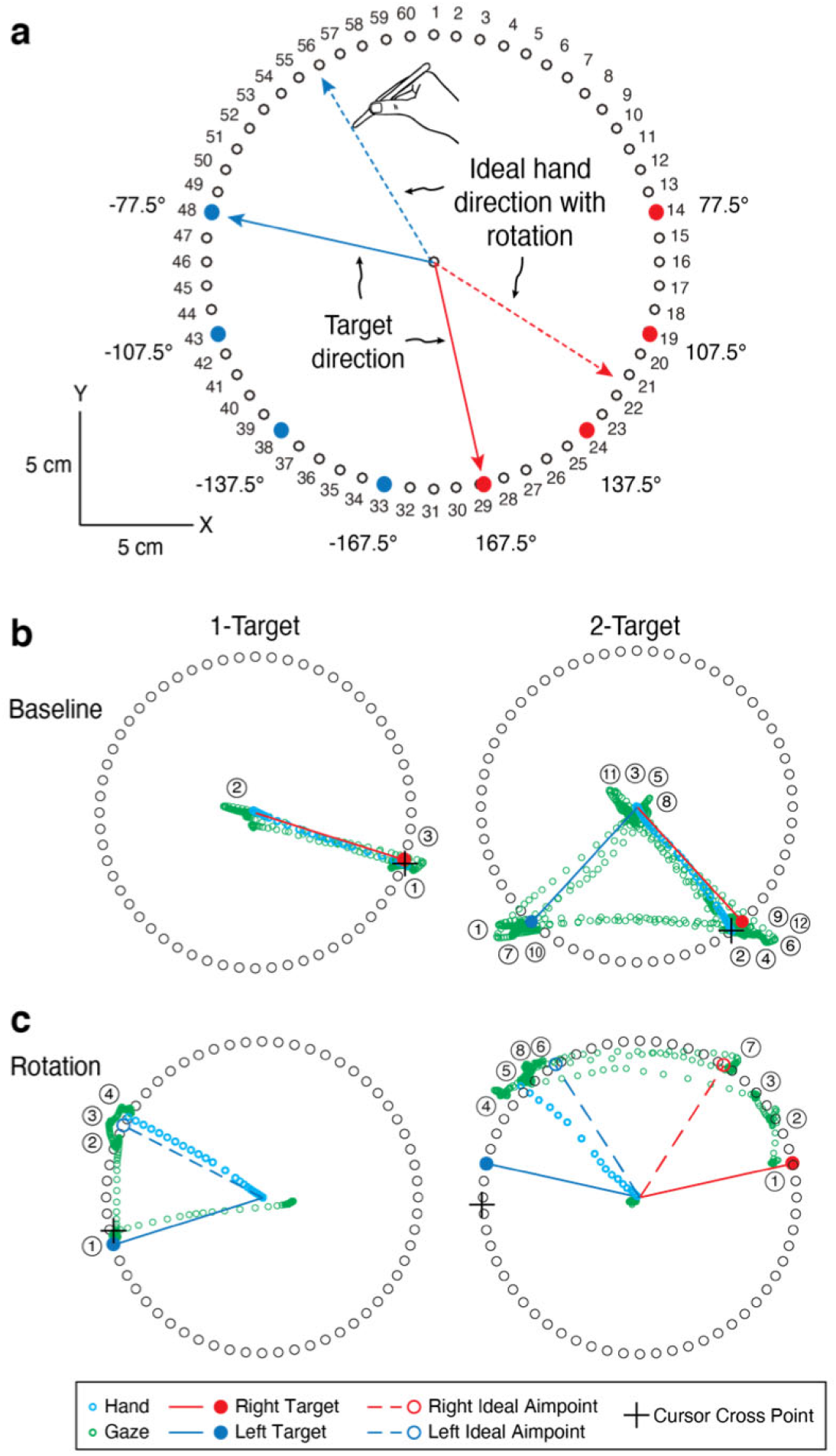
Experimental paradigm and illustrative trials. a, Participants moved the tip of a hand-held stylus across a horizontal surface to move a cursor from central start position to a visual target presented on a vertical monitor. In 1-target trials, either a blue or a red target was displayed and, in 2-target trials, one blue and one red target were displayed. Targets were displayed on a visible ring composed of 60 small circles. The filled blue and red circles indicate the 4 possible locations of the blue and red targets on the left and right sides of the ring, respectively. After a preview period, either the single target, or one of the two potential targets was cued, providing the go signal for the reach. In the rotation phase of the experiment, visuomotor rotations were applied, requiring the participant to move the stylus in a direction rotated 45° CW or CCW from the blue or red target, respectively, to bring the cursor to the target. In Report-and-Reach trials, the circles were numbered and the participant had to indicate the number of the circle they intended to reach toward before executing the reach. b, Gaze paths (green circles) and hand paths (cyan circles) from illustrative 1-target and 2-target reach trials in the baseline phase (when no rotations were applied). The paths are shown from the time of target presentation to the time the cursor crossed the ring boundary. The circled numbers indicate successive fixations. c, Gaze and hand paths from illustrative 1- and 2-target reach trials in the rotation phase (when rotations were applied).

In the ‘report’ and ‘rotation’ phases of the experiment, we used visuomotor rotations to decouple the visual goal locations from the corresponding motor goal locations. This key manipulation, wherein the visual feedback of the cursor movement was rotated about the central start position, allowed us to distinguish gaze fixations tied to the location of the visual target(s) versus the location of the motor goal(s). The learning of visuomotor rotations has been shown to reflect the summation of two separate, but interacting components (Miyamoto et al., 2020; Taylor and Ivry, 2011). The explicit component constitutes a re-aiming strategy, wherein the hand is aimed away from the visual target, in the direction opposite of the rotation. This component has been shown to drive a fast change in hand movement direction early in the learning process (de Brouwer et al., 2018; Taylor et al., 2014). The implicit component, by contrast, involves the automatic (i.e., not under voluntary control) adaptation of the mapping between motor commands and their sensory consequences, resulting in gradual changes in hand movement direction during learning. In our task, visual feedback of the cursor was rotated about the hand start position by 45° clockwise (CW) in trials in which the red target was cued (i.e., for rightward movements), and by 45° counter clockwise (CCW) in trials in which the blue target was cued (i.e., for leftward movements). Thus, to successfully hit the target, the participant had to specify motor goal locations, via the explicit component, to move the stylus in a direction rotated 45° CCW or CW, respectively, from the target (see dashed lines in Fig. 1a). We used opposite rotations for the red and blue targets to limit implicit adaptation over the course of the experiment (Herzfeld et al., 2014; Wigmore et al., 2002). In addition, we sought to maximize the explicit component—and thus the separability of the motor goal location from the corresponding visual target location—by informing participants, after the first rotation trial with each of the target colours, that they could counteract the visuomotor rotation by aiming in a different direction than the visual target. To assess the contribution of the explicit component during the task, and provide a basis for interpreting gaze fixations associated with motor goal locations, we measured the magnitude of the explicit component in reach-and-report trials after the introduction of the rotation (see below).

Figure 1b shows gaze (green) and hand (blue) paths in illustrative 1- and 2-target reach trials taken from the baseline phase (with no rotations applied). The paths are shown from the preview period to the time the cursor crossed the ring. The circled numbers indicate successive gaze fixations—the locations of which correspond to dense regions on the green path—following the initial fixation at the start location. In this example 1-target trial, the participant fixated the visual target (fixation 1), then the center start location (fixation 2), and then the target again during the preview period (fixation 3), where it remained during the reach. In the 2-target trial, gaze shifted between the center start location and each potential target, as well as between the two potential targets, before shifting to the cued target (fixation 12), where it remained during the reach.

Figure 1c shows corresponding gaze and hand paths in illustrative 1- and 2-target reach trials taken from the rotation phase. In the 1-target trial, gaze was initially directed to the visual target (fixation 1) and then shifted, over several fixations (fixations 2 to 4), towards the participant’s explicit aimpoint, rotated ~45° CW from the blue target. This gaze behaviour is consistent with previous work using a similar paradigm and demonstrates encoding of the hand movement (i.e., motor) goal, in addition to the visual goal (i.e., the target) prior to movement (de Brouwer et al., 2018). In the 2-target trial, gaze initially shifted to the red target (fixation 1) before shifting (fixations 2 and 3) towards the red aimpoint rotated ~45° CW. (Note that there is no gaze path from the central fixation point to the first fixation because the participant blinked during this gaze shift.) Although participants often fixated both visual targets, in this trial gaze then shifted midway between the blue target and blue aimpoint rotated 45° CCW (fixation 4) before shifting (fixations 5 and 6) toward the blue aimpoint. Gaze then shifted between the two aimpoints (fixations 7 and 8) suggesting that the motor goal locations associated with each potential target were being held in memory (see below) until the blue target was cued.

### Reaching and Reporting Behaviour

Figure 2a shows, for a representative participant, the hand angle, relative to the cued target, at the moment the cursor crossed the ring, as a function of trial number in the baseline, report, rotation, and no feedback phase of the experiment. The symbols show individual trials, and the vertical blue and red lines show separate averages for blue and red targets, respectively. In the baseline phase, containing 64 intermixed 1- and 2-target trials, reaches were directed to the cued target (i.e., errors are distributed around 0°).

**Figure 2.**
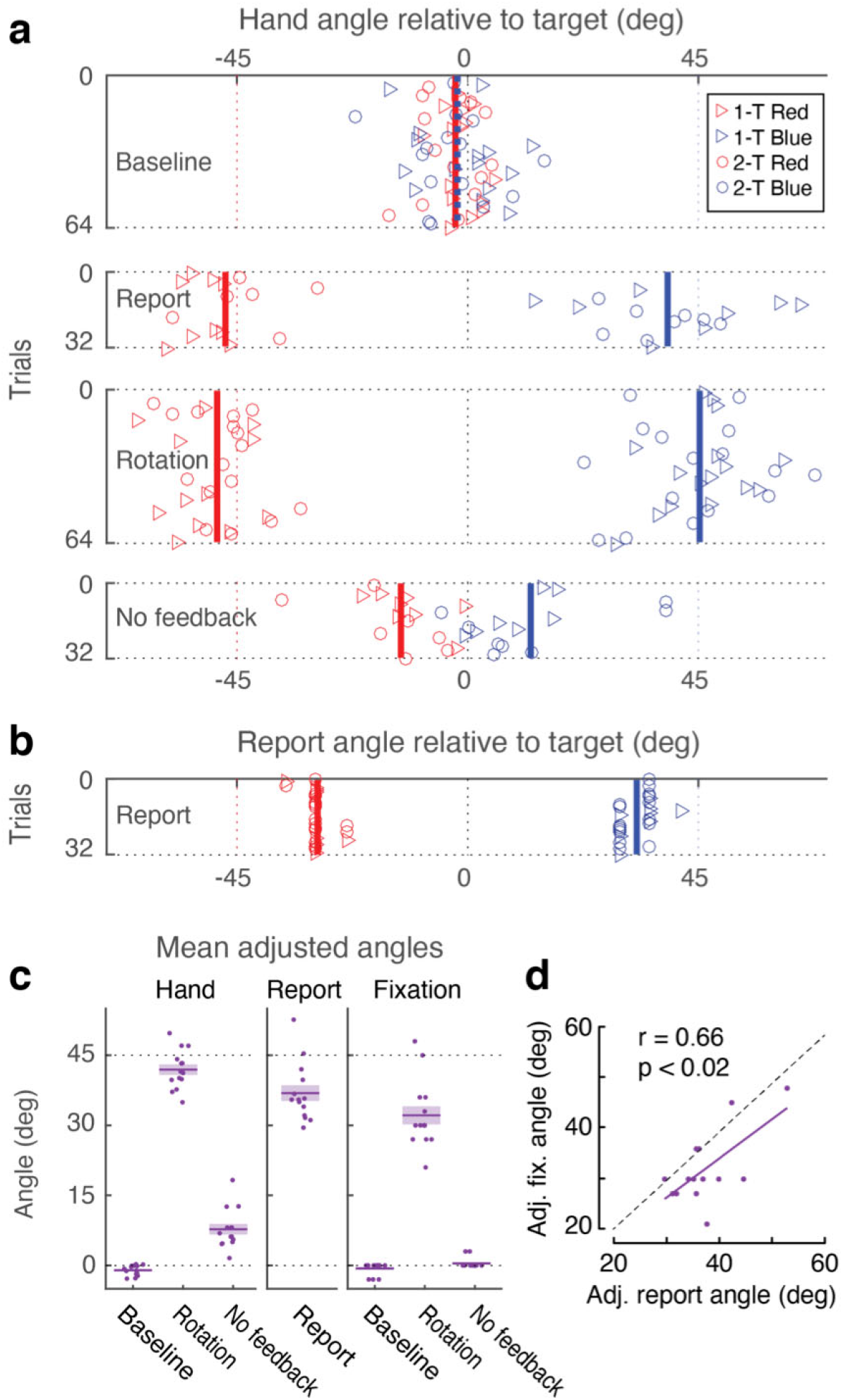
Reach and reporting behaviour of a representative participant (a, b), and across participants (c, d). a, Hand angle, relative to the cued target, at the moment the cursor reached the ring, in each trial in the four successive phases of the experiment. Symbols show individual trials with the colour indicating the cued target and the shape indicating the number of targets displayed. Vertical red and blue lines show the average across trials in which the red or blue target was cued, respectively. b, Verbally reported intended aiming direction in report-and-reach trials during the report phase of the experiment. Symbols and lines as in a. c, To combine the data across the opposing rotations, we computed adjusted hand, report, and fixation angles by negating the angles for the red targets. The left panel shows the mean adjusted hand angle, based on participant averages (dots), in the baseline, rotation, and no feedback phases. The middle panel shows the mean adjusted verbally reported angle in the report phase. The right panel shows the mean adjusted fixation angle, based on participant modes (dots), in the baseline, rotation, and no feedback phases. Short horizontal purple bars and shading show the group mean ±1 standard error. d, Linear relationship between adjusted fixation angle and adjusted report angle. Dots represent individual participants; the purple line shows the best linear fit to the participant data.

Upon completion of the baseline phase, visuomotor rotations of −45° and +45° were applied during reaches to the blue and red targets, respectively. To measure and facilitate the implementation of an explicit re-aiming strategy, participants then completed a series of report-and-reach trials (Taylor et al., 2014). In these trials, the numbers 1 to 60 were displayed next to the circles of the ring (Fig. 1a). In 1-target report-and-reach trials, participants were asked to report the number of the circle they intended to aim towards in order to move the cursor to the target. In 2-target report-and-reach trials, they were asked to report, for each potential target, the number of the circle they intended to reach towards if that target were selected (i.e, they reported two numbers). After the participant reported the target number(s), one target was cued and the participant executed the reach to that target. Participants completed a block of eight 1-target report-and-reach trials for each target color, followed by 32 intermixed red- and blue-cued 1- and 2-target report-and-reach trials. Our analysis of the report phase focused on the latter 32 trials, where the explicit component has largely stabilized.

As illustrated in Figure 2a (Report), the representative participant successfully counteracted the rotation by moving their hand approximately +45° and −45° away from the blue and red targets, respectively (thus moving the cursor to the target). Figure 2b shows that the aimpoints verbally reported by this participant were rotated approximately ±35° from the target, indicating that they primarily counteracted the visuomotor rotation through the use of an explicit re-aiming strategy, with the remainder (approximately ±10°) being achieved through implicit adaptation (Taylor et al., 2014).

Following the report-and-reach trials, participants completed 64 reach trials with the visuomotor rotations applied, but without the numbering of the ring of circles and the reporting procedure. As illustrated in Figure 2a (Rotation), the representative participant successfully moved the cursor to the targets by reaching +45° and −45° away from the blue and red targets, respectively.

After the rotation phase, participants completed an additional 32 reach trials in which visual feedback of the cursor was removed (no feedback phase). In these trials, participants were told that the rotation was now turned off and were instructed to reach directly to the target when it was cued, allowing for the measurement of implicit adaptation (Taylor et al., 2014). As shown in Figure 2a (No feedback), this implicit component (or after-effect) was approximately ±10° in our representative participant. This is consistent with the observation that the reported aiming angle was ±35°, summing up to a hand angle of ±45°, and indicates that the magnitude of the explicit component was preserved throughout the rotation phase.

To combine the data across the two opposing rotations, we computed, for each participant, adjusted hand and report angles by negating the angles for the red targets. Figure 2c shows the mean adjusted hand angle, based on participant averages, in the baseline, rotation, and no feedback phases, as well as the mean adjusted verbally reported angle in the report phase. The report angle (M = 36.9°; SE = 1.7°) and the aftereffect in the no feedback phase at the end of the experiment (M = 7.8°; SE = 1.1°) summed to approximately 45°, as shown above for our representative participant. This finding indicates that the magnitude of the explicit component remained consistent throughout the 64 trials of the rotation phase. (Note that the right side of Fig. 2c and Fig. 2d, which describes gaze behaviour during the task, will be described below.)

### Spatial Distributions of Fixations

Figure 3a shows, for the same representative participant shown in Figure 2a, the angle, relative to the closest target, of all fixations during the target preview period as a function of trial number and phase. (Note that there were often multiple fixations in a given trial.) As illustrated in the figure, during both baseline and no feedback reach trials, fixations were directed close to the visual target(s). During the rotation phase, however, a large proportion of fixations—in both 1-target (triangles) and 2-target (circles) trials—were directed close to the reported aimpoints that this participant had previously reported in the report phase (red and blue vertical lines, also shown in Fig. 2b). Figure 3b shows, for the same participant, the distribution of fixation time as a percentage of total fixation time (including central fixations) during the preview period, at each fixation angle (6° bins), for both the baseline and rotation phases. Separate distributions are shown for 1-target and 2-target trials and for fixations closest to the blue and red targets. It is clear that in the baseline phase, this participant spent most time fixating near the visual target, while in the rotation phase some time was spent near the visual target, but most time was spent fixating near the ‘aimpoint’. To combine the data across the two potential targets, we computed the adjusted fixation angle by negating the angle for the red target. Figure 3c shows, for both phases, the percentage of preview fixation time as a function of adjusted fixation angle, with separate distributions shown for 1- and 2-target trials. Figure 3d shows the distribution of preview fixation time as a function of adjusted fixation angle, averaged across participants. Whereas a single peak at the target (0°) was observed in the baseline phase, two separate peaks, one at the visual target and one in the vicinity of the aimpoint, were observed in the rotation phase.

**Figure 3.**
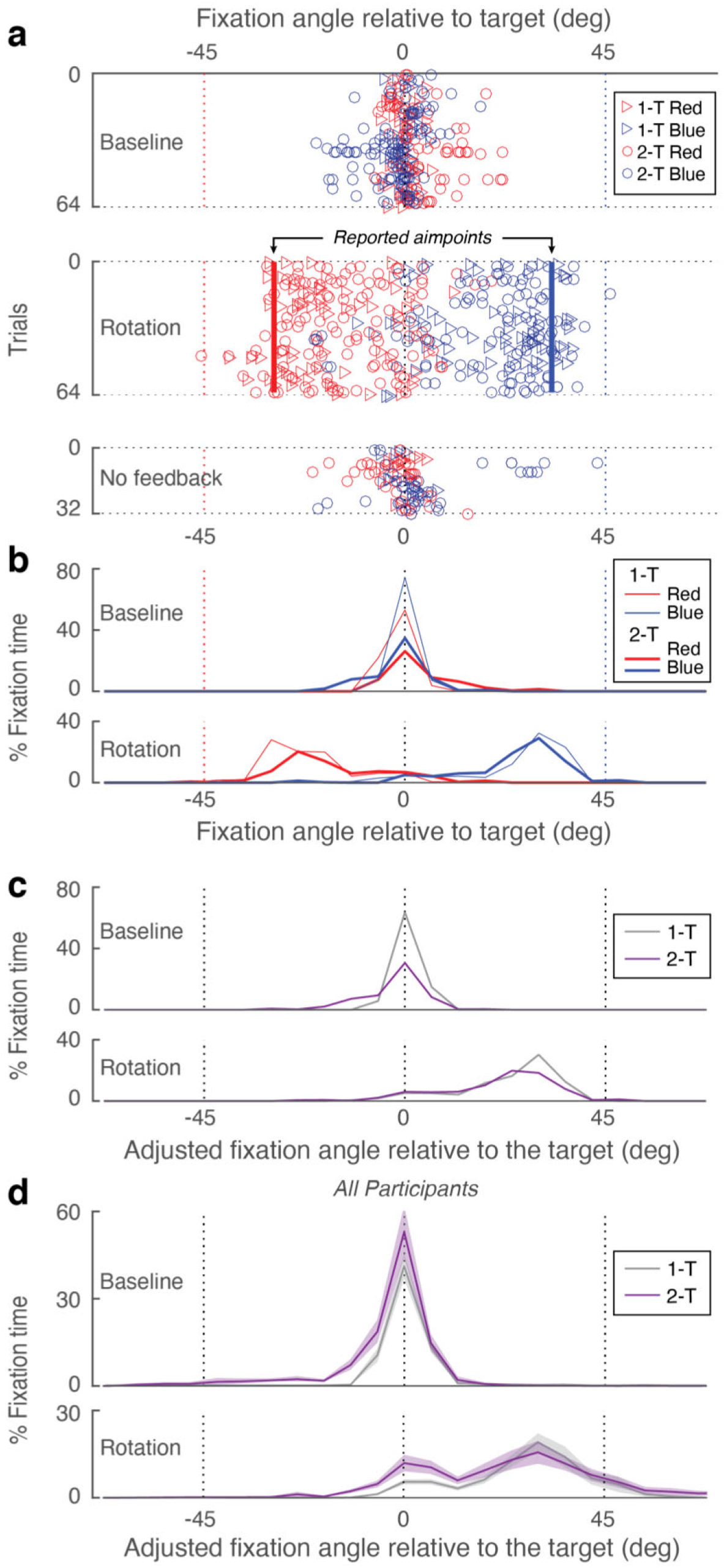
Gaze behaviour. a-c, Results from the same representative participant shown in Fig. 2. a, Angle, relative to the closest target, of all fixations during the preview period. Symbols show individual trials with the colour and shape indicating the closest potential target and the number of targets displayed, respectively. b, Percentage of total fixation time during the preview period in the baseline and rotation phases as a function of fixation angle (6° bins), with separate distributions shown for 1- and 2-target trials and for fixations assigned to the blue and red targets. c, Percentage of preview fixation time in the baseline and rotation phases as a function of adjusted fixation angle (computed by negating the angle for the red targets), with separate distributions shown for 1- and 2-target trials. Data combined across blue and red targets. d, Means, averaged across participants, of the distributions shown in c. Height of the shaded regions represents ±1 standard error.

Next, we determined the location of the peak (or mode) of the fixation time distribution for each participant and phase (baseline, rotation and no feedback), combining 1- and 2-target trials. These modes, along with the mean across participants, are shown in the right panel of Figure 2c to allow for direct comparison with the reaching and report data (discussed above) over the same phases of the task. Note that the mean modal fixation angle during the rotation phase (M = 31.7°; SE = 2.2°) was slightly smaller than the mean reported aimpoint in the report phase (M = 36.9°, SE = 1.7°). This difference is likely due to the fact that our analysis considers all gaze fixations during the preview period and that gaze tended to shift, over two or more fixations, from the target towards the aimpoint (as illustrated in Fig. 1c). Importantly, across participants, the mean fixation angle during the rotation phase correlated with the mean verbally reported aimpoint during the Report phase (Fig. 2d). Together, these findings indicate that participants’ gaze behaviour provides a good covert indicator of their explicit re-aiming strategy, and thus the specification of motor goals prior to target selection.

### Time Course of Within Trial Fixations

To investigate the temporal pattern of gaze fixations during target preview and reach execution of trials in the rotation phase, we defined three spatial zones: a central zone, a target zone, and an aimpoint zone (see inset in Fig. 4; note that in 2-target trials, there were two target zones and two aimpoint zones). Figure 4 displays the time-varying probability, within a trial, of fixating within each of the zones, averaged across participants. The initial sequence of fixations was similar in 1- and 2-target trials. Participants typically fixated the central starting point (orange trace) at the beginning of the trial. After about 300 ms, the probability of fixating the target(s) (black trace) increased sharply and, after about a further 200 ms, the probability of fixating the aimpoint(s) (green trace) increased. In 1-target trials, the probability of fixating in the aimpoint zone remained high until the end of the preview phase. In contrast, in 2-target trials the probability of fixating the aimpoint decreased while the probability of fixating the central zone increased, presumably to ‘wait’ until one of the two potential targets was cued. Towards the end of the reaction time interval, and throughout the movement time interval, the probability of fixating the target increased, presumably to verify the landing position of the cursor relative to the target. In summary, although the time course of zone fixations differed somewhat between 1- and 2-target trials, in both types of trials gaze was often directed to the aimpoint zone(s) during the preview period.

**Figure 4.**
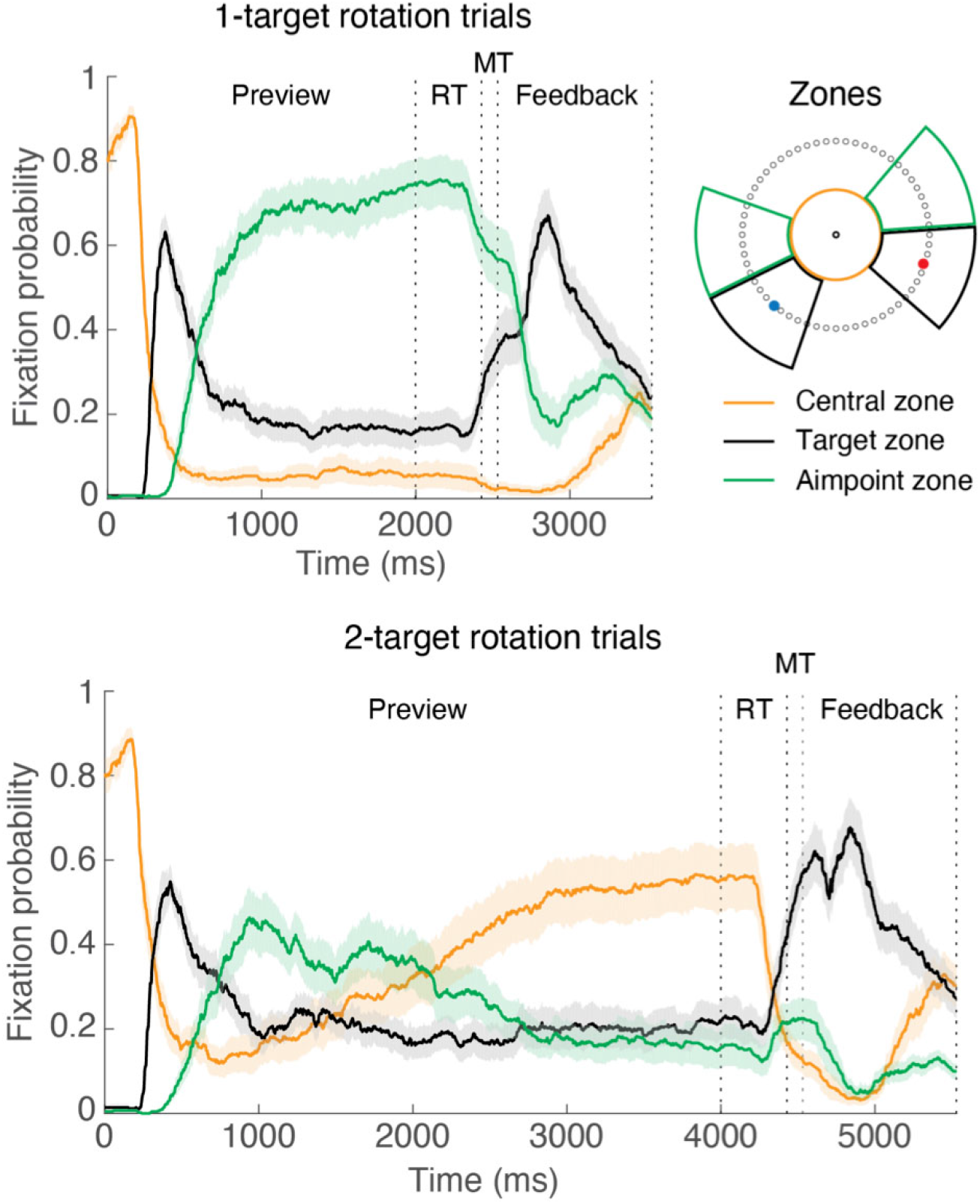
Time course of within trial fixations. Time varying probability of fixating within the target, aimpoint, and central zones in 1-target (top) and 2-target (bottom) trials. The traces represent means, averaged across participants, and the shaded regions represent ± 1 standard error. The reaction time (RT) and movement time (MT) intervals are normalized to the median durations across all included trials, averaged across participants.

### Target and Aimpoint Fixation Probabilities

Having established that fixations observed within the aimpoint zone indicate the specification of motor goals, a key question is how frequently participants fixated aimpoint zones. Figures 5a and b show the frequency with which participants fixated within the target and aimpoint zones in the baseline and rotation phases of the experiment, respectively. Note that these plots express fixation frequency as a percentage of the total number of targets. In the baseline phase (Fig. 5a), participants almost always fixated the target zone in 1-target trials and fixated the majority (M=81%; SE=4%) of target zones in 2-target trials. As expected, participants very rarely fixated the aimpoint zone—as defined for the rotation phase—in the baseline phase, indicating that aimpoint zone fixations during the rotation phase are task-specific. In the rotation phase of the experiment (Fig. 5b), the probability of fixating target zones was a little lower than in the baseline phase zones. However, even in 2-target trials, participants fixated 65% (SE=5%) of the target zones, and thus often fixated both potential targets during the delay period. On average, the frequency with which participants fixated the aimpoint zones (M=57%; SE=7%) was comparable to the frequency with which they fixated the target zones, although there was considerable variability across participants in 2-target trials. Thus, in the rotation phase, aimpoint locations (i.e., motor goals) and visual target location (i.e., visual goals) became similarly salient on average.

**Figure 5.**
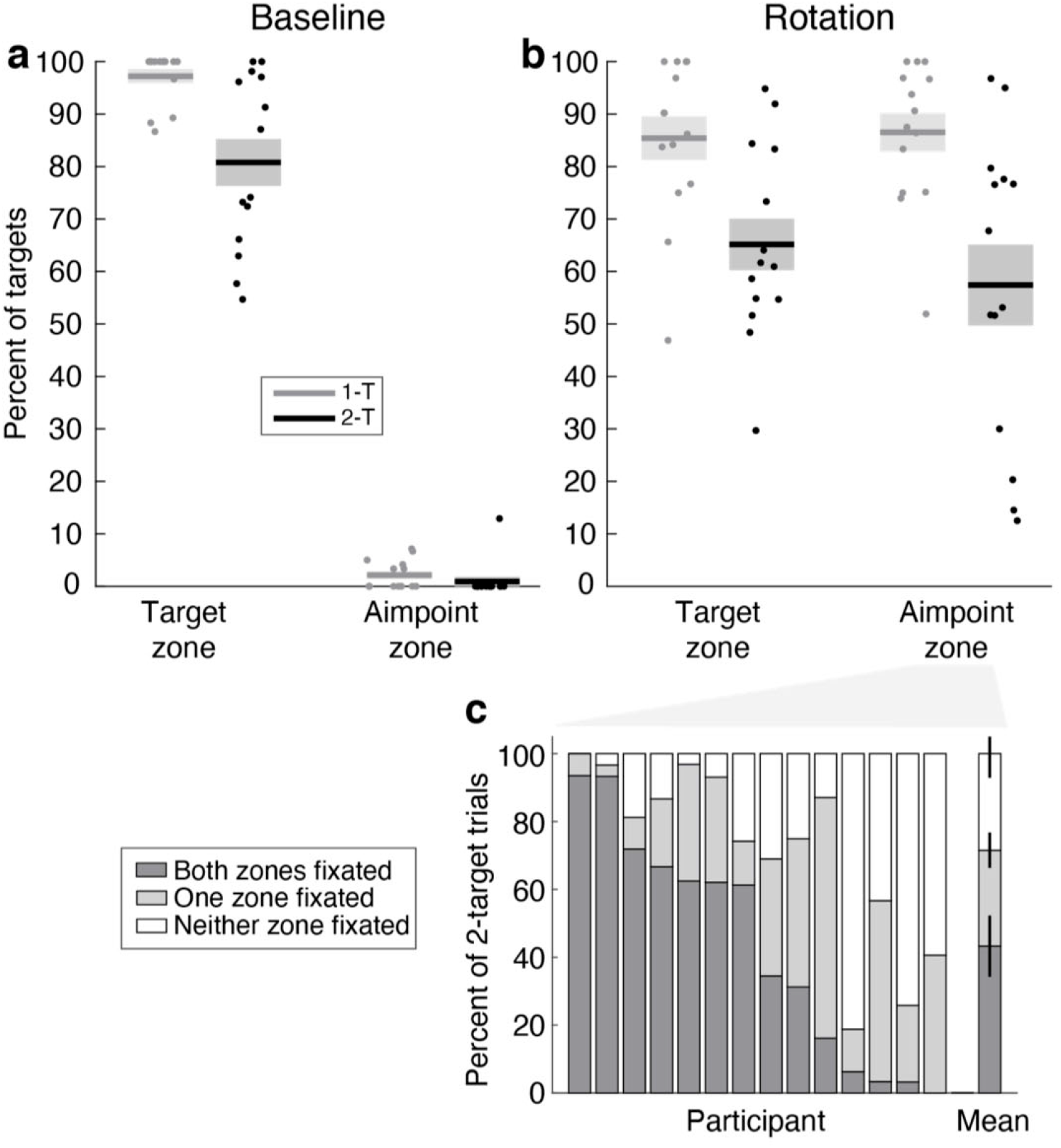
Target and aimpoint fixation probabilities, as a percentage of the total number of targets. a, Probabilities of fixating the target and aimpoint zones in 1-target (1-T) and 2-target (2-T) trials in the baseline phase, expressed as a percentage of the total number of targets in each phase. b, Corresponding percentages for the rotation phase. Each dot represents the data of an individual participant and the horizontal lines and shaded areas depict group means ± 1 standard error of the mean. c, Probability, expressed as a percentage of all 2-target trials, that a fixation occurred in both, one, or none of the aimpoint zones in the rotation phases, for each participant and the average across the group, where the heights of the vertical lines represents ± 1 standard error.

To test whether participants specified single or multiple motor goals prior to target selection, we examined how often, in 2-target trials during the rotation phase, participants fixated both aimpoint zones, only one aimpoint zone, or neither aimpoint zone. Figure 6c shows the probability, for each individual participant as well as for the group, that a fixation occurred in both, one, or none of the aimpoint zones, expressed as a percentage of 2-target trials. We found that the relative frequency of the three aimpoint encoding strategies varied markedly across participants. Thus, half of the participants fixated in both aimpoint zones in the majority (>60%) of 2-target trials, whereas the other half of the participants were more likely to fixate in one or neither of the aimpoint zones. Importantly, most individual participants exhibited a mixture of encoding strategies across trials. This variability both within and across participants challenges that notion that any single model—i.e., parallel specification (e.g., Cisek, 2012), stay-or-switch (Dekleva et al., 2018) or serial—can account for participant behaviour.

**Figure 6.**
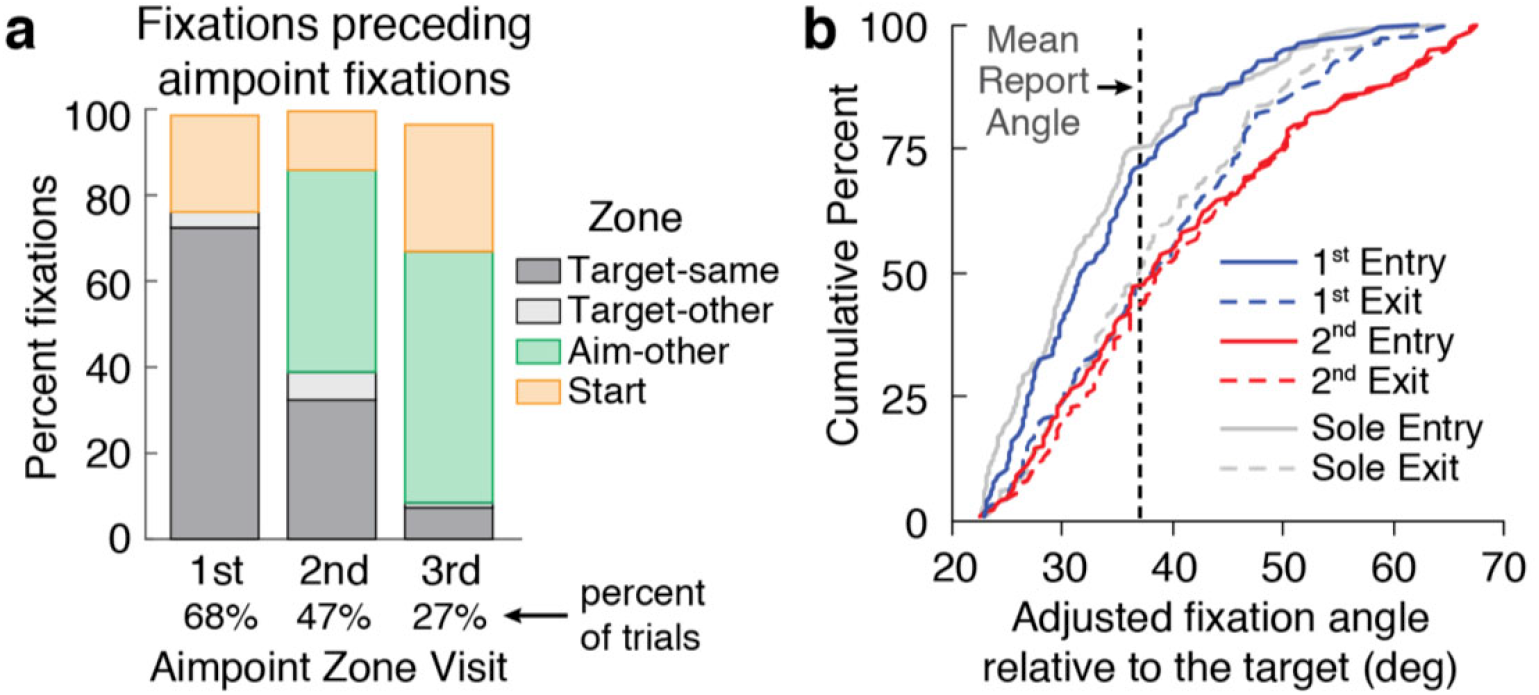
Aimpoint re-visits in 2-target trials. a, Probability that the fixation preceding the first, second, and third gaze visits to a given aimpoint zone was in the corresponding target zone, other target zone, other aimpoint zone, and start zone. b, Cumulative distributions of the entry (solid traces) and exit (dashed traces) fixation angles for the first (blue) and second (red) visits to a given aimpoint zone in instances in which the zone was visited and then revisited after visiting the other aimpoint zone. Corresponding gray traces show distributions of the sole entries and exits of a aimpoint zone in instances in which that zone was visited once.

### Aimpoint Zone Re-Visits

Although fixating both aimpoint zones during the preview period is consistent with the idea that two competing motor goals are specified and then held in memory until the reach is cued, an alternative interpretation is that participants encoded each motor goal sequentially such that the first motor goal is replaced in memory when the participant opts to encode the other motor goal. We found that, during the preview period, gaze often ‘visited’ a given aimpoint zone two, and occasionally three, times, where each visit could involve several fixations. To test whether participants maintained previously encoded motor goals in memory, we carried out two analyses that compared gaze behaviour, in 2 target trials, associated with initial visits and gaze behaviour associated with re-visits.

We first examined the location from which participants launched eye movements to bring gaze into an aimpoint zone. As shown in Fig. 6a, the first aimpoint zone visit (present in 68% of trials) was most often preceded by a fixation in the corresponding visual target zone (72% of cases), suggesting that the motor goal location was derived (and specified immediately) following the visual target location. However, the second and third visits to the aimpoint zone (present in 47 and 27% of trials, respectively) were most often preceded by a fixation in the aimpoint zone for the other target (47 and 58% of cases, respectively). This result suggests that, after participants initially fixated a given aimpoint, they kept that motor goal location in memory such that they could return their gaze directly to it without having to first fixate the corresponding visual target.

The second analysis focused on trials in which gaze visited a given aimpoint zone and then re-visited that zone after visiting the *other* aimpoint zone in between. As noted above, we observed that on the first visit to a given aimpoint zone, the initial or ‘entry’ fixation tended to undershoot the ideal aimpoint, and was followed by one or two additional saccades that brought gaze closer to the ideal aimpoint (see Fig. 1c). This gaze behaviour presumably arises from a stepwise process engaged in determining the aimpoint relative to the visual target. In contrast, we observed that when the aimpoint zone was re-visited, with an intervening visit to the other aimpoint, the entry fixation tended to be close to the ideal aimpoint (also illustrated in Fig. 1c). If substantiated, these observations would suggest that the first motor goal was kept in memory when the second motor goal was encoded, and that, therefore, both motor goals were maintained simultaneously in memory.

To test the hypothesis that participants can hold *two* motor goals in memory, we selected instances (N=132), from 2-target trials in the rotation phase, in which, during the preview period, gaze visited (i.e., entered and exited) a given aimpoint zone and then re-visited that aimpoint zone after having visited the other aimpoint zone. The solid blue trace in Fig. 6b shows the distribution of fixation angles (relative to the target) of the first fixation when gaze visited an aimpoint zone for the first time (1st entry). The dashed blue trace shows the fixation angle of the last fixation before gaze then exited the aimpoint zone for the first time (1st exit). The median fixation angles of the 1st entry and 1st exit were 31.8° and 38.2°, respectively, consistent with the observation that gaze traversed the aimpoint zone during the first visit. Note that the median exit angle is closer to the mean report angle (36.9°; vertical dashed line) than the median entry angle, suggesting that, during the first visit, participants were actively determining the desired aimpoint location.

The solid and dashed red traces show the distributions of fixation angles for the 2nd entry and 2nd exit; i.e., the first and last fixations for the second visit to the same aimpoint zone. In contrast to the first entry and exit angles, both these had medians (38.2 and 38.8°, respectively) which were close to the fixation angle of the 1st exit as well as the mean verbal report angle. This suggests that having visited the aimpoint once, the location was held in memory and used to guide gaze when revisiting the aimpoint zone. Kolmogorov-Smirnov tests revealed that the distributions of entry and exit fixation angles were significantly different (p<0.001) for the first aimpoint zone visit but not the second visit (p=0.29). For completeness, we also examined instances, from 2-target trials in the rotation phase, in which an aimpoint was only entered and exited once (N=168). The solid and dashed gray traces in Fig. 6b show the distributions of these sole entries and exits. The median entry and exit fixation angles were 30.6° and 37.0°, respectively, again indicating that when first visiting an aimpoint zone, gaze tended to arrive short of the actual aimpoint and then traversed the zone towards this aimpoint. These results provide strong evidence that participants stored motor goals in memory after encoding them, and that encoding a second motor goal does not interfere with memory of the first motor goal. Thus, the results indicate that participants could, and often did, maintain two motor goals in memory at the same time.

## Discussion

To investigate how the brain represents competing reach options under target uncertainty, we measured participant’s gaze behaviour while viewing two potential targets, one of which was cued as the reach target after a preview period. Critically, we applied opposing visuomotor rotations to the two targets so that we could dissociate eye fixations related to the visual location of each target from eye fixations related to the motor goal location; i.e., the location that participants aimed their hand toward in order to bring the rotated cursor, controlled by the hand, to the target. We found that, during the preview period, participants generally fixated the visual targets. In terms of motor goal locations, we found that individual participants exhibited a mixture of gaze strategies, across trials, whereby they either fixated both motor goal locations, one of these locations, or neither motor goal location during the preview period. Analysis of gaze behaviour in trials in which gaze re-visited a given motor goal location after visiting the other motor goal location indicated that both motor goals were simultaneously retained in memory during the preview period. These results provide evidence that, at the level of single trials, the brain often encodes multiple motor goals prior to target selection, but may also encode either one or no motor goals.

To date, research examining the encoding of competing targets of action has tended to argue exclusively for one of three models: the parallel specification model (Cisek, 2007; Cisek and Kalaska, 2005; Thura and Cisek, 2014), the stay-or-switch model (Dekleva et al., 2018), and traditional serial models (McClelland, 1979; Sternberg, 1969). In the context of delayed reach tasks in which two potential targets are presented, these models posit that, prior to target selection, motor goals are specified for both potential targets, a single potential target, or neither potential target. Our finding that, across trials, individual participants employ a mixture of encoding strategies—variously specifying two, one or no motor goals prior to target selection—challenges the notion that any single model can account for how the brain represents competing reach options. The choice of strategy at a particular moment in time may depend on a number of factors including attentional and motivational states, the cognitive effort and memory demands involved in specifying motor goals, and real or perceived benefits associated with advance motor goal specification (Cisek and Kalaska, 2010; Gallivan et al., 2016a, 2015; Thura and Cisek, 2014).

It is important to note that the frequency with which aimpoint zones were *fixated* may underestimate the frequency with which motor goals were actually *specified* during the preview period. Whereas an aimpoint zone fixation during the preview period provides evidence that the corresponding motor goal was specified prior to target selection, the absence of an aimpoint fixation does not necessarily imply that the motor goal was not specified using peripheral vision. In our previous study in which only single targets were presented during the preview period (de Brouwer et al., 2018), we observed that participants who fixated the motor goal exhibited fast learning consistent with the implementation of an explicit strategy whereby they specified, and aimed toward, the motor goal. In contrast, most of the participants who did not fixate the motor goal during the preview period exhibited more gradual learning consistent with implicit adaptation without explicit motor goal specification. However, a couple of participants exhibited explicit learning without fixating the aimpoint, indicating that they specified, and aimed toward, the motor goal without fixating it. Thus, we may be underestimating the frequency of trials in which both motor goals were specified.

The idea that participants can flexibly use different encoding strategies has implications for interpretations from previous neurophysiological studies. For example, in their study examining the encoding of potential reach targets, Dekleva and colleagues (2018) simultaneously recorded activity from populations of neurons in dorsal premotor cortex during a delayed response task. They tested how well their data were fit by single and dual target encoding models and concluded that, at the level of single trials, only one or two response options was represented at a time during the delay period. However, their analyses, perhaps necessarily, did not consider the possibility of flexible encoding, which, as we have shown in the current study, may be quite variable across participants.

We found that, in 2-target trials, participants often fixated both aimpoint zones during the preview phase. One interpretation of this gaze behaviour is that both motor goals are encoded and maintained in memory prior to target selection. However, an alternative interpretation is that participants forget the motor goal associated with the previously fixated aimpoint when they fixate the other aimpoint, effectively switching which motor goal is held in memory. Importantly, our analysis of aimpoint zone visits and re-visits provides strong support for the former interpretation. We found that when gaze first visited a given aimpoint zone, this was typically preceded by a fixation of the corresponding target. This suggests that participants actively determined the motor goal location relative to the target location. In contrast, we found that when gaze re-visited a given aimpoint zone, the initial fixation in the zone was most often preceded by a fixation of the other aimpoint. This suggests the motor goal location, specified during the first visit, was kept in memory such that the participant did not have to re-fixate the visual target in order to locate the motor goal.

Critically, we also found that when, during the preview period, gaze re-visited a given aimpoint zone after visiting the other aimpoint zone, the entry fixation was far closer to the verbally reported aimpoint than the entry fixation on the first visit to that aimpoint zone. (The latter tended to undershoot the reported aimpoint and was followed by one or more saccades that brought gaze toward the reported aimpoint.) This result provides strong evidence that participants encoded and remembered motor goal locations, even after subsequently encoding the other motor goal location, and thus could hold two motor goals in memory while waiting for one of the potential reach targets to be cued. We suggest that aimpoint re-visits, which often involved alternating fixations between aimpoints, may be akin to rehearsal strategies that people employ to maintain items in working memory (Baddeley, 2007; Baddeley and Hitch, 1974). From this perspective, re-visits can be viewed as serving to maintain and reinforce multiple motor goal locations in memory.

In visually guided actions, task-specific proactive eye movements are crucial for planning and control (Johansson et al., 2001; Land et al., 1999; Land and McLeod, 2000) and may be viewed as an integral component of the overall motor program for the task (Flanagan et al., 2013; Flanagan and Johansson, 2003; Land and Furneaux, 1997; Rotman et al., 2006). Moreover, information gained through task-specific eye movements need not be used immediately to guide action, but can be buffered for use in guiding forthcoming actions (Land and Furneaux, 1997; Land and Lee, 1994; Land and Tatler, 2009). In our task, a fixation of an aimpoint or motor goal, during the preview period, can thus be viewed as a key component to specifying the potential reaching movement to the corresponding target. In principle, fixating an aimpoint can provide both visual and extra-retinal (i.e., gaze-related proprioceptive or efference copy signals) information about the location of the intended spatial goal of the hand movement that may be required (Prablanc et al., 1986, 1979; Prablanc and Martin, 1992). Note that although the aimpoint was ~10 degrees away from the location to which the hand was directed due to implicit adaptation, there is evidence that gaze-related signals may still be used to guide the hand when fixating a location that is close to the hand’s target (Neggers and Bekkering, 2001). Moreover, it is possible that the processing of gaze-related signals for hand guidance incorporates implicit adaptation.

We have argued that fixating the aimpoint associated with a potential target, during the preview phase, is tantamount to specifying or encoding the motor goal associated with that target. Whereas many authors, including ourselves, have previously suggested that potential reaching movements are ‘planned’ in advance of target selection, we recognize that motor goal specification should not be equated with movement planning. The latter involves a number of components, ranging, depending on the theoretical account, from trajectory specification and optimization (Flash and Hogan, 1985; Harris and Wolpert, 1998) to the setting of feedback gains to optimize feedback control (Scott, 2004; Todorov, 2004; Todorov and Jordan, 2002). We cannot know, based on gaze behaviour, the extent to which such processes are completed, in advance, for each potential action. However, all accounts of movement planning and control involve motor goal specification.

In summary, we have provided evidence based on gaze behaviour in a delayed reaching task with two potential targets, that participants employed a mixture of strategies whereby, across trials, they may specify motor goals for both targets, one target, or neither target prior to target selection. This finding challenges theoretical accounts that have assumed that participants inflexibly use a single encoding strategy when confronted with competing potential targets.

## Methods

### Participants

Fifteen participants (M_age_ = 20.5, SD_age_ = 1.0; 13 women) were recruited from the student population at Queen’s University and provided written informed consent prior to completing the experiment. All participants were right-handed as verified by the Edinburgh Handedness Inventory (Oldfield, 1971) and were compensated $10 CAD for their time. The Queen’s University General Ethics Board approved all experimental procedures. A target sample size of 14-16 participants was specified in advance based on previous studies examining eye movements in action tasks, and our expectation that, if the main experimental effect is present, it should be observed at the single-subject level in nearly all participants. One participant was excluded from all analyses because they rarely moved their gaze from the central starting position during the trial (i.e., both the preview and execute periods), even in the baseline condition. Thus, their gaze behaviour could not be used to examine whether, or not, they prepared movements in advance initiating reaches.

### Apparatus

Participants made center-out reaching movements to visual targets by moving the tip of a hand-held stylus across a horizontal digitizing tablet (active area 31.1 × 21.6 cm; Wacom Intuos PTH-851, Wacom, Kazo, Sataima, Japan). All visual stimuli were presented on a vertical computer monitor (display size 47.5 × 26.5 cm; resolution 1920 × 1080 pixels; refresh rate 60 Hz). The participant’s head was supported by a chin and forehead rest placed ~50 cm in front of the monitor (10 cm on the screen corresponded to ~11.3 degrees of visual angle). The position of the tip of the stylus was sampled at 100 Hz and the participant’s view of their hand was occluded. Movements of the right eye were recorded at 500 Hz using a video-based eye tracker (EyeLink 1000, SR Research Ltd, Kanata, Ontario) located below the computer monitor, following a standard nine-point calibration.

### Stimuli

The position of the tip of the stylus—which we will refer to as the hand position—was represented on the monitor as a circular cursor (1 cm diameter) that moved with a ratio of 1.3 times the displacement of the hand. Sixty open white circles (0.6 cm diameter; 6° spacing) were displayed in a ring (radius 10 cm) around a central starting circle (1 cm diameter). In report-and-reach trials (see below), target numbers (1-60) were displayed eccentric to the targets (see Fig. 1A). Movements were made from the starting circle to targets (2 cm diameter) located on the ring.

Throughout the experiment there were two target colours. Red targets were always presented on the right side of the ring and appeared at one of four locations (77.5, 107.5, 137.5, or 167.5°), whereas blue targets always appeared on the left side of the ring at one of four mirrored locations (−77.5, −107.5, −137.5, or −167.5°), as displayed in Figure 1A. Zero degrees corresponded to up on the screen (y-direction), and positive angles correspond to clockwise rotations. Note that these target locations were selected so that there would be little ambiguity in determining which target (blue or red), or corresponding aimpoint (see below), a given eye fixation was directed towards.

### Procedure

Participants began each trial by moving the cursor to the central start position. Once this position was maintained for 0.5 s, either one unfilled target (red or blue) or two unfilled targets (one red and one blue) appeared on the ring. Following a fixed 2 s (1-target trials) or 4 s (2-target trials) delay period, either the single target or one of the two targets was filled in, providing the go-signal to reach to that target as quickly and accurately as possible. All targets were presented in pseudorandom order such that the same (combination of) location(s) did not appear on two consecutive trials.

Participants were instructed to make a quick movement that “sliced” through the target. Visual feedback of the cursor was provided throughout the movement and, when the cursor crossed the ring, a circle (equal in size to the size of the cursor) was drawn at the crossing location to provide additional feedback about reach accuracy. Participants earned points for hitting the target, provided they initiated their movement between 100 and 600 ms following the go-signal. If the participant anticipated the go-signal (i.e., initiated movement less than 100 ms after the go-signal) or took longer than 600 ms to initiate the movement, the message “too early” or “too late” was displayed, respectively, and the trial was aborted. A trial was considered a hit if any part of the cursor contacted any part of the target. The message “hit” or “miss” was displayed in all trials that met the reaction time criteria.

Participants first completed 64 reach trials with veridical cursor feedback (i.e., no visuomotor rotations were applied; baseline phase). This phase included 32 1-target trials (16 red and 16 blue) and 32 2-target trials (16 red target cued and 16 blue target cued) presented in a pseudorandom order with all target locations cued an equal number of times.

Following the baseline phase, participants performed the report phase in which visuomotor rotations were applied. Specifically, visual feedback of the cursor was rotated about the hand start position, +45° in trials in which the red target was cued and −45° in trials in which the blue target was cued. Participants first completed a single reach trial with a single target, after which the experimenter informed participants that they would have to counteract a visuomotor rotation to successfully hit the target, encouraging participants to implement a re-aiming strategy. Opposing visuomotor rotations were used for the red and blue targets to guard against implicit adaptation to the rotations (Wigmore et al., 2002). To measure the magnitude of the re-aiming strategy, participants performed report-and-reach trials in which the target numbers were displayed and they were asked to verbally report the number of the circle they intended to reach towards (Taylor et al., 2014). In 1-target report-and-reach trials, participants reported a single number, and in 2-target trials they were asked to report a number for each target (red then blue). After this report was completed, either the single target, or one of the two targets, was filled in, providing the cue to initiate a reach (as in reach trials). Participants first completed a block of 8 1-target report-and-reach trials with the red targets and a block of 8 1-target report-and-reach trials with the blue targets, being informed about the rotation after the first trial of each target color. Participants then completed a block of randomly intermixed 1-target and 2-target report-and reach trials, consisting of 16 1-target (8 red and 8 blue) and 16 2-target trials (8 red target cued and 8 blue target cued) with all target locations cued an equal number of times (32 trials in total). Following the report phase, participants completed the rotation phase, which consisted of reach trials without report. This phase was identical to the baseline phase, with the exception that visual feedback of the cursor was rotated by ±45°. After completing the rotation phase, participants performed a phase without visual feedback (no feedback phase), which allowed us to assess the contributions of implicit and explicit learning. Participants were told that the rotation was turned off and instructed to aim directly at the target (Morehead et al., 2017). This phase involved 16 1-target (8 red and 8 blue) and 16 2-target (8 red cued and 8 blue cued) trials that were presented in a pseudorandom order and target locations were cued an equal number of times (32 trials in total). Participants were given 30 s breaks between blocks of trials and additional breaks halfway through the two 64 trial blocks experienced during the baseline and rotation phases.

### Data Analysis

For all analyses, we only included trials where movements were initiated within 100 and 600 ms following the go-signal and the cursor crossed the ring within 400 ms after movement onset (93% of trials). To obtain a measure of task performance, we used the endpoint hand angle relative to the target angle at the moment the cursor crossed the ring. Explicit learning (i.e., the magnitude of re-aiming) was quantified by converting the verbally reported landmark number to an angle relative to the target angle. For each participant, we computed mean hand and explicit angles for each phase and target, and then averaged values across the blue and red target after mirroring the angles for the red target across the vertical midline.

We analysed gaze data for reach trials without report, including all trials in which there were no blinks or missing data for at least 50% of the time from initial target presentation until the cursor crossed the ring (91% of reach without report trials). Blinks were first removed from the *x* and *y* gaze positions and these signals were then low-pass filtered using a second-order recursive Butterworth filter with a cut-off frequency of 50 Hz. The filtered *x* and *y* gaze positions were used to calculate horizontal, vertical, and resultant gaze velocity. Data were drift-corrected offline by computing the median *x* and *y* gaze position at target onset (when gaze is still at the start position) across all trials, for each block separately, and shifting the data by aligning the median *x* and *y* gaze positions to the start position. Next, the onset and offset of saccades were defined based on resultant gaze velocity with saccades identified as having a resultant velocity above 200 mm/s (or ~22.6 °/s) for five or more consecutive samples (i.e., 10 ms). Onsets were defined as the last of five samples below the threshold of 200 mm/s and offsets were defined as the first of five samples below this threshold. We only considered saccades with a minimum displacement of 5 mm. Fixations were then defined as periods of 50 or more consecutive samples (100 ms) during which neither a blink nor a saccade occurred. For each fixation, we computed the mean *x* and *y* position.

The resulting fixation positions were used to quantify 1) distributions of fixation positions, 2) the time course of gaze over a single trial in the rotation phase, and 3) the probability of fixating targets and aimpoints. For the first analysis, we first computed the angle relative to each target for all fixations within 50 and 150% of the target distance (i.e., non-central fixations). Fixations were binned into 60 bins, with the center of the bins corresponding to the angles of the open circles forming the ring, and the widths of the bins corresponding to the angular distance between two adjacent circles (i.e., 6°). We computed the fixation time in each bin relative to the target as a percentage of the total fixation time (including central fixations, excluding blinks and saccades) during the target preview of each trial. For 2-target trials, fixations were assigned to the closest target, and the fixation time in each bin was computed as a percentage of half of the total fixation time. For each participant, we computed the distribution of percentage fixation time in each bin for the blue and red target and for 1- and 2-target trials separately. We then computed the combined distribution, including all valid trials and mirroring fixation angles for the red target, for the baseline phase and in the rotation phase. The modes of the combined distribution were taken as a measure of the fixation angle. For 12 out of the 14 participants, this value was close to the ‘ideal aimpoint’ in the rotation phase. For 2 participants, we manually selected the second highest peak of the distribution as a measure of fixation angle, since the highest peak occurred at 0° (i.e., at the target).

For the second and third analyses, we defined fixation zones: a central zone, a target zone, and an aimpoint zone (see inset in Fig. 4). The central zone was a circle, centered on the hand start position, with a radius of 50% of the distance to the target ring. Target and aimpoint zones were defined for each target; i.e., there were two target zones and two aimpoint zones in two target trials. These zones were 45° wide wedges between 50 and 150% of the distance to the target ring. The target zone was centered on the target, and the aimpoint zone was centered 45° CW or CCW from the target, depending on whether the target was red or blue. That is, the aimpoint zone was centered on the hand location required to bring the cursor to the target when the visuomotor rotations were applied. To examine gaze patterns over the time course of a single trial in the rotation phase, we computed the probability of fixation in each of the zones for each time sample. The probabilities were computed after normalizing the reaction time interval and reach interval (i.e., movement time interval) of each trial to the mean duration of that interval across all participants, for 1-target trials and 2-target trials separately.

In the third analysis, we assessed how often participants fixated the targets and aimpoints. We computed the probability of a fixation in each zone during the baseline and rotation phase, for 1- and 2-target trials separately, as a percentage of the total number of targets. To assess whether participants prepared a movement to both targets in 2-target trials, we computed the probability that a fixation occurred in both, one, or neither of the target and aimpoint zones.

Finally, we examined the temporal pattern of fixations in the target and aimpoint zones in two-target trials. Specifically, we expected that the first fixation in the aimpoint zone would be preceded by a fixation in the target zone on the same side of the display, while later aimpoint zone fixations would be preceded increasingly often by an aimpoint zone fixation on the opposite side of the display. This would allow participants to keep both aimpoint locations in memory in anticipation of one of the targets being cued. To test this, we computed the probability of fixation in each of the target zones, the aimpoint zone for the other target, and the start zone, separating fixations preceding the first, second and third aimpoint fixation. When sequential fixations in the same aimpoint zone occurred, we only used the first of these fixations. In addition, we examined the fixation angles of the first (entry) and last (exit) fixations for each gaze visit to a given aimpoint zone, during which there could be several fixations. We first selected all instances, in 2-target trials, in which an aimpoint zone was visited more than once, mirroring the angles for the red target so that all angles were positive. Next, we obtained the angle of the entry and exit fixations for the first and second visits of the aimpoint zone. We then tested whether the fixation angle changes during a visit to the aimpoint zone, i.e., by making small saccades within the aimpoint zone.

## Acknowledgements

This work was supported by a Natural Sciences and Engineering Research Council (NSERC) Discovery grant awarded to J.R.F., an operating grant from the Canadian Institutes of Health Research (CIHR) awarded to J.R.F. and J.P.G. The authors would like to thank Martin York and Sean Hickman for technical assistance.

## Author Contributions

All authors contributed to the design of the experiment and writing the paper. M.J.C. and L.S. performed the research. A.J.d.B and J.R.F. analysed and interpreted the data.

## References

Baddeley A. 2007. Working Memory, Thought, and Action. doi:10.1093/acprof:oso/9780198528012.001.0001

Baddeley AD, Hitch G. 1974. Working Memory. Psychology of Learning and Motivation. doi:10.1016/s0079-7421(08)60452-1

Chapman CS, Gallivan JP, Wood DK, Milne JL, Culham JC, Goodale MA. 2010. Reaching for the unknown: multiple target encoding and real-time decision-making in a rapid reach task. Cognition 116:168–176.

Christopoulos V, Bonaiuto J, Andersen RA. 2015. A biologically plausible computational theory for value integration and action selection in decisions with competing alternatives. PLoS Comput Biol 11:e1004104.

Christopoulos V, Schrater PR. 2015. Dynamic Integration of Value Information into a Common Probability Currency as a Theory for Flexible Decision Making. PLoS Comput Biol 11:e1004402.

Cisek P. 2012. Making decisions through a distributed consensus. Curr Opin Neurobiol 22:927–936.

Cisek P. 2007. Cortical mechanisms of action selection: the affordance competition hypothesis. Philos Trans R Soc Lond B Biol Sci 362:1585–1599.

Cisek P, Kalaska JF. 2010. Neural mechanisms for interacting with a world full of action choices. Annu Rev Neurosci 33:269–298.

Cisek P, Kalaska JF. 2005. Neural correlates of reaching decisions in dorsal premotor cortex: specification of multiple direction choices and final selection of action. Neuron 45:801–814.

Coallier É, Michelet T, Kalaska JF. 2015. Dorsal premotor cortex: neural correlates of reach target decisions based on a color-location matching rule and conflicting sensory evidence. J Neurophysiol 113:3543–3573.

Crawford JD, Medendorp WP, Marotta JJ. 2004. Spatial transformations for eye-hand coordination. J Neurophysiol 92:10–19.

de Brouwer AJ, Albaghdadi M, Flanagan JR, Gallivan JP. 2018. Using gaze behavior to parcellate the explicit and implicit contributions to visuomotor learning. J Neurophysiol 120:1602–1615.

Dekleva BM, Kording KP, Miller LE. 2018. Single reach plans in dorsal premotor cortex during a two-target task. Nat Commun 9:3556.

Flanagan JR, Johansson RS. 2003. Action plans used in action observation. Nature 424:769–771.

Flanagan JR, Rotman G, Reichelt AF, Johansson RS. 2013. The role of observers’ gaze behaviour when watching object manipulation tasks: predicting and evaluating the consequences of action. Philos Trans R Soc Lond B Biol Sci 368:20130063.

Flash T, Hogan N. 1985. The coordination of arm movements: an experimentally confirmed mathematical model. J Neurosci 5:1688–1703.

Gallivan JP, Barton KS, Chapman CS, Wolpert DM, Flanagan JR. 2015. Action plan co-optimization reveals the parallel encoding of competing reach movements. Nat Commun 6:7428.

Gallivan JP, Bowman NAR, Chapman CS, Wolpert DM, Flanagan JR. 2016a. The sequential encoding of competing action goals involves dynamic restructuring of motor plans in working memory. J Neurophysiol 115:3113–3122.

Gallivan JP, Chapman CS, Wolpert DM, Flanagan JR. 2018. Decision-making in sensorimotor control. Nat Rev Neurosci 19:519–534.

Gallivan JP, Chapman CS, Wood DK, Milne JL, Ansari D, Culham JC, Goodale MA. 2011. One to four, and nothing more: nonconscious parallel individuation of objects during action planning. Psychol Sci 22:803–811.

Gallivan JP, Logan L, Wolpert DM, Flanagan JR. 2016b. Parallel specification of competing sensorimotor control policies for alternative action options. Nat Neurosci 19:320–326.

Gallivan JP, Stewart BM, Baugh LA, Wolpert DM, Flanagan JR. 2017. Rapid Automatic Motor Encoding of Competing Reach Options. Cell Rep 18:1619–1626.

Haith AM, Huberdeau DM, Krakauer JW. 2015. Hedging your bets: intermediate movements as optimal behavior in the context of an incomplete decision. PLoS Comput Biol 11:e1004171.

Haith AM, Pakpoor J, Krakauer JW. 2016. Independence of Movement Preparation and Movement Initiation. J Neurosci 36:3007–3015.

Harris CM, Wolpert DM. 1998. Signal-dependent noise determines motor planning. Nature 394:780–784.

Herzfeld DJ, Vaswani PA, Marko MK, Shadmehr R. 2014. A memory of errors in sensorimotor learning. Science 345:1349–1353.

Hudson TE, Maloney LT, Landy MS. 2007. Movement planning with probabilistic target information. J Neurophysiol 98:3034–3046.

Johansson RS, Westling G, Bäckström A, Flanagan JR. 2001. Eye-hand coordination in object manipulation. J Neurosci 21:6917–6932.

Klaes C, Westendorff S, Chakrabarti S, Gail A. 2011. Choosing Goals, Not Rules: Deciding among Rule-Based Action Plans. Neuron 70:536–548.

Land MF, Furneaux S. 1997. The knowledge base of the oculomotor system. Philos Trans R Soc Lond B Biol Sci 352:1231–1239.

Land MF, Lee DN. 1994. Where we look when we steer. Nature. doi:10.1038/369742a0

Land MF, McLeod P. 2000. From eye movements to actions: how batsmen hit the ball. Nat Neurosci 3:1340–1345.

Land M, Mennie N, Rusted J. 1999. The roles of vision and eye movements in the control of activities of daily living. Perception 28:1311–1328.

Land M, Tatler B. 2009. Looking and Acting: Vision and Eye Movements in Natural Behaviour. Oxford University Press.

McClelland JL. 1979. On the time relations of mental processes: An examination of systems of processes in cascade. Psychol Rev 86:287–330.

Miyamoto YR, Wang S, Smith MA. 2020. Implicit adaptation compensates for erratic explicit strategy in human motor learning. Nature Neuroscience. doi:10.1038/s41593-020-0600-3

Morehead JR, Taylor JA, Parvin DE, Ivry RB. 2017. Characteristics of Implicit Sensorimotor Adaptation Revealed by Task-irrelevant Clamped Feedback. J Cogn Neurosci 29:1061–1074.

Nashed JY, Diamond JS, Gallivan JP, Wolpert DM, Flanagan JR. 2017. Grip force when reaching with target uncertainty provides evidence for motor optimization over averaging. Sci Rep 7:11703.

Neggers SF, Bekkering H. 2001. Gaze anchoring to a pointing target is present during the entire pointing movement and is driven by a non-visual signal. J Neurophysiol 86:961–970.

Oldfield RC. 1971. The assessment and analysis of handedness: the Edinburgh inventory. Neuropsychologia 9:97–113.

Pastor-Bernier A, Cisek P. 2011. Neural correlates of biased competition in premotor cortex. J Neurosci 31:7083–7088.

Prablanc C, Echallier JE, Jeannerod M, Komilis E. 1979. Optimal response of eye and hand motor systems in pointing at a visual target. II. Static and dynamic visual cues in the control of hand movement. Biol Cybern 35:183–187.

Prablanc C, Martin O. 1992. Automatic control during hand reaching at undetected two-dimensional target displacements. J Neurophysiol 67:455–469.

Prablanc C, Pélisson D, Goodale MA. 1986. Visual control of reaching movements without vision of the limb. I. Role of retinal feedback of target position in guiding the hand. Exp Brain Res 62:293–302.

Rand MK, Rentsch S. 2015. Gaze locations affect explicit process but not implicit process during visuomotor adaptation. J Neurophysiol 113:88–99.

Rentsch S, Rand MK. 2014. Eye-hand coordination during visuomotor adaptation with different rotation angles. PLoS One 9:e109819.

Rotman G, Troje NF, Johansson RS, Flanagan JR. 2006. Eye movements when observing predictable and unpredictable actions. J Neurophysiol 96:1358–1369.

Scott SH. 2004. Optimal feedback control and the neural basis of volitional motor control. Nat Rev Neurosci 5:532–546.

Sternberg S. 1969. Memory-scanning: mental processes revealed by reaction-time experiments. Am Sci 57:421–457.

Stewart BM, Baugh LA, Gallivan JP, Flanagan JR. 2013. Simultaneous encoding of the direction and orientation of potential targets during reach planning: evidence of multiple competing reach plans. J Neurophysiol 110:807–816.

Stewart BM, Gallivan JP, Baugh LA, Flanagan JR. 2014. Motor, not visual, encoding of potential reach targets. Curr Biol 24:R953–4.

Suriya-Arunroj L, Gail A. 2019. Complementary encoding of priors in monkey frontoparietal network supports a dual process of decision-making. Elife 8. doi:10.7554/eLife.47581

Taylor JA, Ivry RB. 2011. Flexible cognitive strategies during motor learning. PLoS Comput Biol 7:e1001096.

Taylor JA, Krakauer JW, Ivry RB. 2014. Explicit and implicit contributions to learning in a sensorimotor adaptation task. J Neurosci 34:3023–3032.

Thura D, Cisek P. 2014. Deliberation and commitment in the premotor and primary motor cortex during dynamic decision making. Neuron 81:1401–1416.

Todorov E. 2004. Optimality principles in sensorimotor control. Nat Neurosci 7:907–915.

Todorov E, Jordan MI. 2002. Optimal feedback control as a theory of motor coordination. Nat Neurosci 5:1226–1235.

Wigmore V, Tong C, Flanagan JR. 2002. Visuomotor rotations of varying size and direction compete for a single internal model in motor working memory. J Exp Psychol Hum Percept Perform 28:447–457.

Wong AL, Haith AM. 2017. Motor planning flexibly optimizes performance under uncertainty about task goals. Nat Commun 8:14624.

